# Zebrafish Xenografts Reveal a Context-dependent Role of PFKFB4 in Melanoma Cell

**DOI:** 10.1101/2025.09.06.674616

**Authors:** Chenxi Zhou, Francesca Lorenzini, Boris Bardot, Vincent Kappès, Laura Fontenille, Karima Kissa, Anne H. Monsoro-Burq

## Abstract

Cutaneous melanoma is a highly metastatic cancer in which tumor cell plasticity drives transitions between proliferative and invasive states. The glycolytic regulator PFKFB4 has been implicated in cancer cell motility, but its role in melanoma progression *in vivo* remains unclear. Using zebrafish larval xenografts, we found that PFKFB4 depletion reduced invasion in MeWo cells without affecting tumor growth, whereas in A375P cells it decreased tumor growth but had no effect on invasion or metastasis. In MeWo, PFKFB4 loss was associated with reduced SNAIL2 expression, while no effect on EMT transcription factors was observed in A375P cells. Rescue experiments indicated that PFKFB4 and SNAIL2 acted independently in regulating Mewo migration *in vitro*. These findings identify PFKFB4 as a context-dependent regulator of melanoma progression.

## INTRODUCTION

Melanoma is the most fatal skin cancer due to its high rate of metastasis^1^. Non-canonical regulators of melanoma metastasis have been recently identified, but how they interact with established mechanisms remains unclear. Being exposed to solar ultraviolet light, malignant skin melanoma has the highest mutation burden across 30 human cancers^2^, and can be subdivided into four subtypes based on their genomic mutation: about 52% of patients harbor a BRAF somatic mutation, around 28% a N/H/K-RAS mutation; 14% present an NF1 mutation. The remaining “triple-negative” ones often present aberrant copy-number of other oncogenes^3^. Several signaling pathways are thus deregulated and promote melanoma growth, invasion, and metastasis. The most common BRAF^V600E^ mutation activates MAPK/ERK pathway^4,5^. RAS activation and PTEN mutations regulate downstream MAPK/ERK as well as PI3K/AKT pathways^3,6^. Alongside this genomic classification, cutaneous melanoma transcriptomics profiles have identified inter- and intra-tumor heterogeneity representing different melanoma phenotypic states, which indicate a balance between a “proliferative” and an “invasive” phenotype^7–9^. The “proliferative” state is also referred to as “melanocytic” due to high expression of MITF and melanocyte markers compared to low MITF expression in the “invasive” state, which is also coupled with dedifferentiation of the cells^10–13^. Single-cell sequencing has revealed additional intermediate states in between those two phenotypes, including “starved” melanoma cells and “neural crest-like” melanoma cells^10,12,14^.

Skin melanoma metastasis is largely driven by melanoma cell plasticity and involves a phenotype switch between a highly proliferative immotile status and a poorly proliferative/highly motile invasive status with decreased E-cadherin and increased N-cadherin levels^15,16^. At the colonization sites, metastatic melanoma cells regain a proliferative status, allowing secondary tumor growth^7,17^. The phenotype switch is controlled by several signaling and molecular pathways, including MITF^7^, MAPK/ERK signaling^18^, inflammatory signaling by TGFβ^8^, and TNFα^19^. Although melanocytes are not epithelial cells, their tumor progression involves an epithelial-to-mesenchymal (EMT)-like process regulated by the main EMT transcription factors (EMT-TFs) SNAIL1/2, TWIST, ZEB1/2, and PRRX1^10,16^. Unlike in other contexts in which they often promote cancer cell metastasis^20^, in melanoma ZEB2 and SNAIL2 mark proliferating melanoma cells located superficially (melanoma in situ), as well as normal melanocytes and nevi^21,22^. High ZEB2 expression in melanoma is associated with high MITF and SOX10 levels, indicating a melanocytic status^22^. Moreover, ZEB2 knockdown blocks melanoma cell proliferation^22^. Similarly, SNAIL2 is highly expressed both in normal skin and proliferative melanoma cells^23^. However, *in vitro*, SNAIL2 overexpression increases N-cadherin levels and activates melanoma cell migration^23^. In contrast, ZEB1 and TWIST, expressed during melanoma vertical growth phase and in metastases *in vivo*, promote invasion *in vitro*^21,24^.

Melanoma phenotype switch is also affected by metabolic rewiring in cancer induced by hypoxia and low glucose tumor microenvironment^25,26^. The rewiring of metabolism could help the cells to adapt to different environments during the metastasis process ^17^. Compared to cancer cells prior to EMT, the cells post-EMT prefer enhanced glycolysis, named the Warburg effect^27,28^. Glycolysis can help to relieve the oxidative stress induced by the high adenosine triphosphate (ATP) production and consumption during cell migration^29,30^. A divergence from glycolysis to pentose phosphate pathway (PPP) is another way to resist oxidative stress^31,32^. This metabolism rewiring relies, in particular, on regulations by 6-phosphofructo-2-kinase/fructose-2,6-biphosphatases enzymes (PFK-2/FBPase-2, PFKFB1-4)^33^.

PFKFB are bifunctional enzymes that have kinase activity transforming fructose-6-phosphate (F-6-P) into fructose-2,6-biphosphate (F-2,6-BP), the most potent positive allosteric effector of phosphofrucokinase-1 (PFK-1), the glycolysis rate-regulating enzyme^34^. By this means, PFKFB controls the glycolysis rate. The phosphatase activity, on the contrary, catalyzes the degradation of F-2,6-BP to F-6-P, which is the entry substrate for PPP. Among the four PFKFB1-4, PFKFB3 and 4 are found to be highly expressed in tumors, including melanoma^35–42^. So far, most studies have addressed PFKFB4 functions as a metabolism selector between glycolysis and PPP metabolism, balancing the energy supply and oxidative homeostasis in cancer^33,42–47^. However, a few studies recently reported novel PFKFB4 non-canonical functions. For example, PFKFB4 promotes breast cancer metastasis either by binding to SRC-3 and phosphorylating it on Ser857 through its kinase activity^48^, or by activating p38 and ERK signaling^49^. Notably, we recently found that PFKFB4 regulates cell migration as a non-canonical AKT regulator, in embryonic neural crest as well as in melanoma^37,50,51^. In melanoma, we found that PFKFB4 interacts with ICMT, a posttranslational modifier of RAS. PFKFB4 promotes ICMT/RAS interaction, controls RAS localization at the plasma membrane, activates AKT signaling, and enhances cell migration *in vitro* independently of glycolysis^37^.

However, it remained unknown, if PFKFB4 also affected cancer cell invasion and metastasis ability *in vivo*, when cells face a diverse and complex environments. Moreover, considering that PFKFB4 regulates AKT, that AKT/GSK3 signaling can modulate SNAIL2 expression and the important functions of EMT-TFs in melanoma plasticity^52–54^, it was important to assess whether PFKFB4 regulation of melanoma migration was dependent on AKT-EMT-TFs activity. In this study, we tested whether PFKFB4 regulated melanoma cell line invasion *in vivo* using a zebrafish xenograft model. We showed that PFKFB4 depletion reduced invasion in MeWo but not in A375P cells, pointing to a cell line-dependent effect. In MeWo cells, loss of PFKFB4 was associated with decreased SNAIL2 expression, while no effect on EMT transcription factors was observed in A375 cells, which express SNAIL2 at very low levels. Taken together, our findings suggest that the regulation of invasion by PFKFB4 is context-dependent and cannot be solely explained by EMT-TF modulation, highlighting the need to explore additional mechanisms.

## MATERIALS AND METHODS

### Patient cohorts and transcriptomic datasets

Melanoma patient cohorts from TCGA (SKCM) and from GSE7553, GSE8401, and GSE3189 datasets were analyzed to assess the relationship between PFKFB4 expression levels and 5-year patient survival (Figure 1), as well as to compare PFKFB4 transcript levels during melanoma progression (Figure S1). Differential PFKFB4 expression in normal skin, nevi, primary melanoma, and metastatic melanoma samples from microarray datasets was evaluated using one-way ANOVA on the R2 Genomic Analysis and Visualization Platform (http://r2.amc.nl). RNA-seq TPM values for the TCGA SKCM cohort were downloaded from UCSC Xena (https://xenabrowser.net). PFKFB4 TPM levels were compared between primary and metastatic tumors using a *t*-test. For survival analysis, patients with metastatic melanoma were stratified into three groups based on PFKFB4 expression: top quartile (high), middle two quartiles (medium), and bottom quartile (low). Kaplan–Meier survival curves were generated using GraphPad Prism.

**Figure 1.**
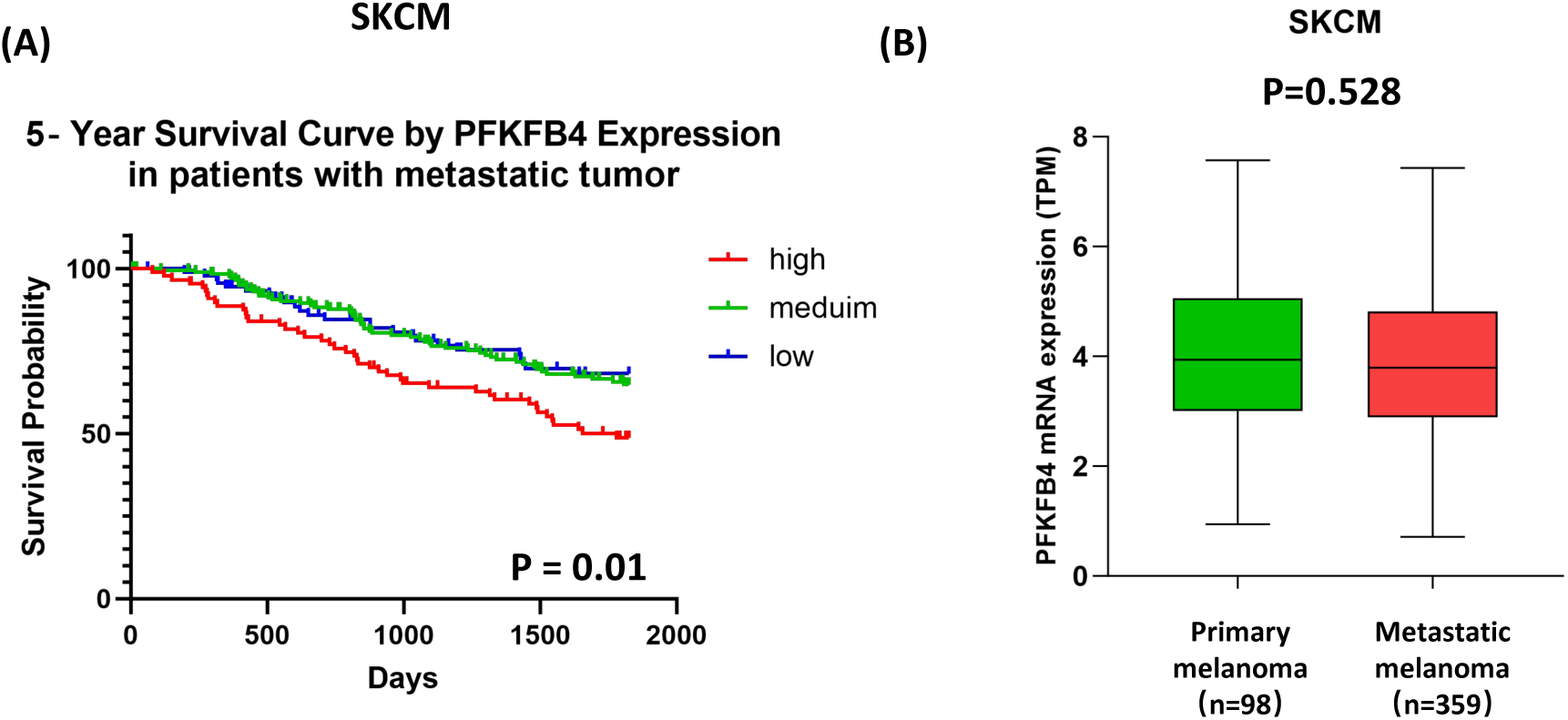
High PFKFB4 expression is associated with reduced survival in metastatic cutaneous melanoma. **(A)** Kaplan–Meier survival curves showing the correlation between PFKFB4 mRNA expression and overall survival in patients with metastatic melanoma. Patients were stratified into three groups based on PFKFB4 expression: top quartile (high), middle two quartiles (medium), and bottom quartile (low). **(B)** Boxplot comparing PFKFB4 expression levels between primary and metastatic melanoma samples. Data were obtained from the TCGA SKCM dataset. Statistical significance was assessed using Mantel-Cox **(A)** and Student’s *t*-test **(B)**.

### Cell culture and transfection

Human metastatic melanoma cell lines MeWo (ATCC #HTB-65)^55^, A375M^55^, and A375P-GFP^56^ (ATCC #CRL-3224), as well as HEK-293FT cells (Thermo Fisher), were maintained in complete RPMI-1640 medium (MeWo, A375M; Gibco, #21875-034) or in complete DMEM (A375P, HEK-293FT; Gibco, # 41965-039), supplemented with 10% fetal bovine serum (Eurobio, #CVFSVF00-0U) and 1% penicillin/streptomycin (Sigma, #P0781).

Twenty-four hours before transfection, cells were seeded in 6-well plates at a density of 2 × 10^5^ cells/well (A375M, A375P) or 3 × 10^5^ cells/well (MeWo). siRNAs (Dharmacon SMARTpool) were transfected at 90 nM using Lipofectamine RNAiMAX (Invitrogen, #2218549). The siRNA sequences targeting human ***PFKFB4*** are: 5′-GAGCGACCATCTTTAATTT-3′, 5′-CATCGTATATTACCTCATG-3′, 5′-GAAATGACCTACGAGGAAA-3′, and 5′-GGGACAGGCCTCAGAACGT-3′. The siRNA sequences targeting human ***SNAI2*** are: 5′-GGACACACAUACAGUGAUU-3′, 5′-UAAAUACUGUGACAAGGAA-3′, 5′-GAAUGUCUCUCCUGCACAA-3′, and 5′-GAAUCUGGCUGCUGUGUAG-3′.

For plasmid transfection, pcDNA, pcDNA-xPFKFB4^37^, or pcDNA-xSNAI2^57^ were introduced at 1 μg/well using Transporter 5 (Polysciences, #26008) according to the manufacturer’s instructions. Cells were harvested and analyzed 48 h post-transfection by Western blotting or RT-qPCR.

### Generation of a stable MeWo-GFP cell line by lentiviral transduction

HEK-293FT cells were cultured in T75 flasks until ∼50% confluence. One hour prior to transfection, culture medium was replaced with fresh medium. For transfection, 10 μg pTRIP-2A-GFP, 10 µg pCMVΔ8.91, 5 µg pVSV-G, and 125 μl 1 M CaCl₂ (Sigma Ultra, C5080) were diluted in H₂O to a final volume of 500 μl. This solution was then added dropwise to 0.5 ml of 2× Hepes-buffered saline (Biochemica Fluka, 51558) and vortexed briefly. The mixture was incubated for 20 min at room temperature (RT) and subsequently added to HEK-293FT cells. After 24 h, the medium was replaced with fresh medium. At 48 h post-transfection, GFP-expressing lentiviral particles were collected by centrifugation at 2500 rpm for 10 min, aliquoted, and stored at –80 °C.

For infection, MeWo cells were seeded in 6-well plates and grown to ∼70% confluence. Cells were then incubated with 1 ml of lentiviral stock per well for 24 h, after which the medium was replaced with fresh medium. Infection efficiency was monitored by GFP fluorescence.

### *In vitro* two-dimensional random cell migration assay

Twenty-four hours after transfection, cells were detached and seeded at 1.5 × 10^4^ cells/well (MeWo) or 1.0 × 10^4^ cells/well (A375P) in 24-well plates pre-coated with fibronectin (40 ng/µl; Sigma, #F1141) and 0.1% bovine serum albumin (BSA). After 24h of culture, cells were subjected to two-dimensional (2D) random migration assays. Nuclei were labeled with the SPY-DNA-555 probe (Spirochrome, SC201) diluted 1:2000, added 4 h prior to image acquisition.

Cell migration was monitored by time-lapse video microscopy (Leica MM AF) equipped with a cell culture chamber (37 °C, humidified atmosphere, 5% CO₂) under phase-contrast and mCherry fluorescence at 10× magnification. Images were acquired every 8 min over a 16 h period using a CoolSnap CCD camera (Photometrics) and Metamorph software (Molecular Devices), yielding a total of 121 frames per field. For each cell line, five independent fields were analyzed. Image sequences were converted to movies using FIJI (https://imagej.nih.gov/ij/). Migration trajectories were obtained either manually, using the Manual Tracking plugin, or automatically. For automated analysis, nuclei were segmented with StarDist 2D^58^, and trajectories were extracted with the TrackMate plugin. Mean cell migration velocities were compared using the Wilcoxon rank-sum test in R v3.6.

### Animal Rearing

Zebrafish (*Danio rerio*) strains were raised and maintained in the AZELEAD Fish Facility under standard conditions^59^. Wildtype adults were crossed to obtain embryos for xenotransplantation procedures. Embryos were initially maintained at 28 °C in fish water supplemented with 0.0002% methylene blue (Sigma, Burlington, MA, USA) as an antifungal agent. After 24 h, embryos were transferred to fish water containing 200 µM Phenylthiourea (PTU) to inhibit pigmentation and enable fluorescent imaging acquisition. All experimental procedures were conducted in accordance with the European guidelines for animal protection and regulations of the French Ministry of Health (authorization F341725).

### Xenograft of human melanoma cells in zebrafish larvae

A375P-GFP and MeWo-GFP cells were harvested 24h after transfection, resuspended in PBS, and injected into the swim bladder of zebrafish larvae at 3 days post fertilization (3 dpf). Injections were performed under a stereomicroscope (Leica M80 Stereo zoom microscope) using a micromanipulator (Narishige, London, UK) and an oil-driven manual microinjector (Cell Tram Vario, Eppendorf, Hamburg, Germany). Larvae were imaged at the day of injection (0 dpi) and at 3 dpi to assess tumor cell dissemination. Imaging was performed on live anesthetized larvae (0.16 mg/mL PBS/Tricaine (MS-222)) using a Zeiss CD7 microscope (ZEISS), acquiring with 52 Z slices per larva. Larvae that already displayed detached at 0 dpi and larvae with an abnormally inflated swim bladder at 3 dpi were excluded from the final analysis.

Detached tumor foci were counted slice by slice using the FIJI cell counter plug-in, with manual labelling of the swim bladder. Tumor volumes were reconstructed and measured with the 3D project and 3D object counter functions in FIJI. Migration distance was defined as the shortest Euclidean distance between the detached foci and the primary tumor boundary. Briefly, (i), the tumor boundary was segmented by Otsu thresholding in FIJI, and its coordinates saved in the ROI manager; (ii), detached foci were manually selected and their coordinates saved in the ROI manager;(iii), Euclidean distances between foci and all tumor boundary coordinates were calculated, and the shortest value was taken as the migration distance. Statistical analysis were performed using unpaired *t-*test in GraphPad Prism.

### Protein extraction, quantification, and western blotting

Forty-eight hours after transfection, cells were washed with PBS and lysed in RIPA buffer (10mM Tris-HCl pH 8, 150 mM NaCl, 1% NP-40, 0,1% SDS, 0,5M sodium deoxycholate), supplemented with protease inhibitors (Roche, # 04693116001) and phosphatase inhibitors (Roche, #04906837001) on ice. Lysates were centrifuged at 16,900 × g for 12 min, and supernatant were collected. Protein concentration was determined using the Bradford reagent (Biorad, #500-0006) by measuring absorbance at 590 nm. Samples were supplemented with 20% Laemli loading buffer and 33mM dithiothreitol (DTT), aliquoted and stored at −20 ℃.

Proteins were denatured at 95 ℃ for 5 min and separated on SDS-PAGE (4% stacking and 12% resolving gels) and transferred onto PVDF membrane (Merck #IPVH00010). Membranes were blocked with 5% milk in TBS containing 0.1% Tween (TBST) for 1 hour at room temperature (RT), then incubated overnight at 4°C with the following primary antibodies diluted in blocking buffer: anti-PFKFB4 (1/2000, abcam #ab137785), anti-ACTIN (1/1000, sigma #A2066), anti-VINCULIN (1/5000, sigma #V9131), anti-SNAIL1 (1/500, Cell signaling #3895), anti-SNAIL2 (1/500, Cell signaling #9585), anti-TWIST (1/500, SANTA CRUZ #sc81417), anti-ZEB1 (1/500, invitrogen #14-9741-82), and anti-ZEB2 (1/1000, Invitrogen #PA5-20980). After five washes with TBST (5 minutes each), membranes were incubated with HRP-conjugated goat anti-rabbit or anti-mouse secondary antibodies (1:20,000) for 1 h at RT. Chemiluminescent signals were detected using a Viber imaging system and quantified using FIJI software.

### RNA extraction and RT-qPCR

Cells were lyzed in TRIzol (Ambion #15596018), and total RNA was extracted using the RNeasy Mini kit (Qiagen #74104) according to the manufacturer’s protocol. Reverse transcription was performed with 1μg RNA at 42 ℃ for 30 min in a reaction containing 0.25μl M-MLV reverse transcriptase (Promega M1701), M-MLV buffer, 0.5μl RNasin (Promega N2111), random hexamers, dNTPs and 1μl 20 mM DTT. Quantitative PCR (qPCR) was carried out using SYBR Green Supermix (Bio-Rad).

The primer sequences used were as follows:

SNAIL1: Fwd 5’-AAGATGCACATCCGAAGCC-3’; Rev5’-CGCAGGTTGGAGCGGTCAGC-3’ SNAIL2: Fwd 5’-ATACCACAACCAGAGATCCTC-3’; Rev 5’GACTCACTCGCCCCAAAGATG3’ TWIST: Fwd 5’-GGGCCGGAGACCTAGATG-3’; Rev 5’-TTTCCAAGAAAATCTTTGGCATA-3’ ZEB2: Fwd 5’-AACAACGAGATTCTACAAGCC-3’; Rev 5’-TCGCGTTCCTCCAGTTTTCTT-3’ **RESULTS**

### PFKFB4 promotes MeWo melanoma cell invasion in a zebrafish xenograft model

We previously showed that PFKFB4 promoted active cell migration in several melanoma cell lines *in vitro*^37^. To assess whether *PFKFB4* expression levels were associated with clinical outcome, we analyzed multiple public melanoma datasets. In the TCGA human cutaneous melanoma cohort, *PFKFB4* mRNA expression was comparable between primary and metastatic tumors, but elevated expression correlated with poorer 5-year survival rate in patients with metastatic melanoma (**Figure 1**). A similar association was previously reported in the GSE65904 microarray dataset^60^. Using the R2genomic Analysis and Visualization Platform (http://r2.amc.nl), we next examined three additional datasets (GSE3189, GSE7553, and GSE8401; **Figure S1. A-C**) which likewise showed no significant difference in *PFKFB4* transcript levels between primary and metastatic melanoma. In GSE7553 (**Figure S1. A**), *PFKFB4* expression appeared higher in metastatic melanoma samples compared with normal skin or melanoma in situ. However, this pattern was not observed in GSE3189 dataset (**Figure S1. B**) and given the very limited number of normal skin samples in both datasets, no firm conclusion can be drawn.

These findings suggested that although *PFKFB4* expression might not reliably distinguish metastatic from primary melanomas at the transcriptomic level, elevated *PFKFB4* expression was associated with poor prognosis and might contribute functionally to melanoma progression. To further explore the role of PFKFB4 in melanoma progression, we employed a larval zebrafish xenograft model^56^ to assess its impact on local invasion and distant metastasis *in vivo*.

The zebrafish swim bladder is an epithelial organ isolated from blood vessels and other tissues, that can serve as a confined site for the initial engraftment of fluorescent tumor cells. During dissemination *in vivo*, the tumor cells may invade surrounding tissues and organs and subsequently metastasize by intravasating through the aorta and spreading via the circulation to distant sites, such as the head or the caudal hematopoietic tissue in the tail^56,61^. By tracking detached melanoma cells in developing larva, we monitored both local invasion and distant metastasis *in vivo* (**Figure 2. A**). For these experiments, we used an A375P-GFP melanoma cell line^56,62,63^ and MeWo cells^37^ newly engineered to stably express GFP. MeWo-GFP cells retained the morphology of parental cells (**Figure S2. A**). The migratory capacity of these GFP-positive cell lines was first assessed *in vitro* using random cell migration assays, where individual cells were tracked for 16 hours by time-lapse microscopy. Consistent with observations in parental cell lines^37^, GFP-positive melanoma cells displayed reduced velocity following PFKFB4 knockdown (**Figure 2. B**). After confirming efficient PFKFB4 depletion by Western blot (**Figure S2. B**), MeWo-GFP or A375P-GFP cells transfected with control siRNA (siCtrl) or PFKFB4-targeting siRNA (siPFKFB4) were injected into the swim bladder of zebrafish larvae at 3 days post-fertilization (3 dpf). Images were acquired at 0 and 3 days post-injection (0 dpi and 3 dpi), and only larvae with no cells outside the swim bladder at 0 dpi and with a normally inflated swim bladder at 3 dpi were included in the analysis (**Figure 2. A**, **Figure S2. C, D**).

**Figure 2.**
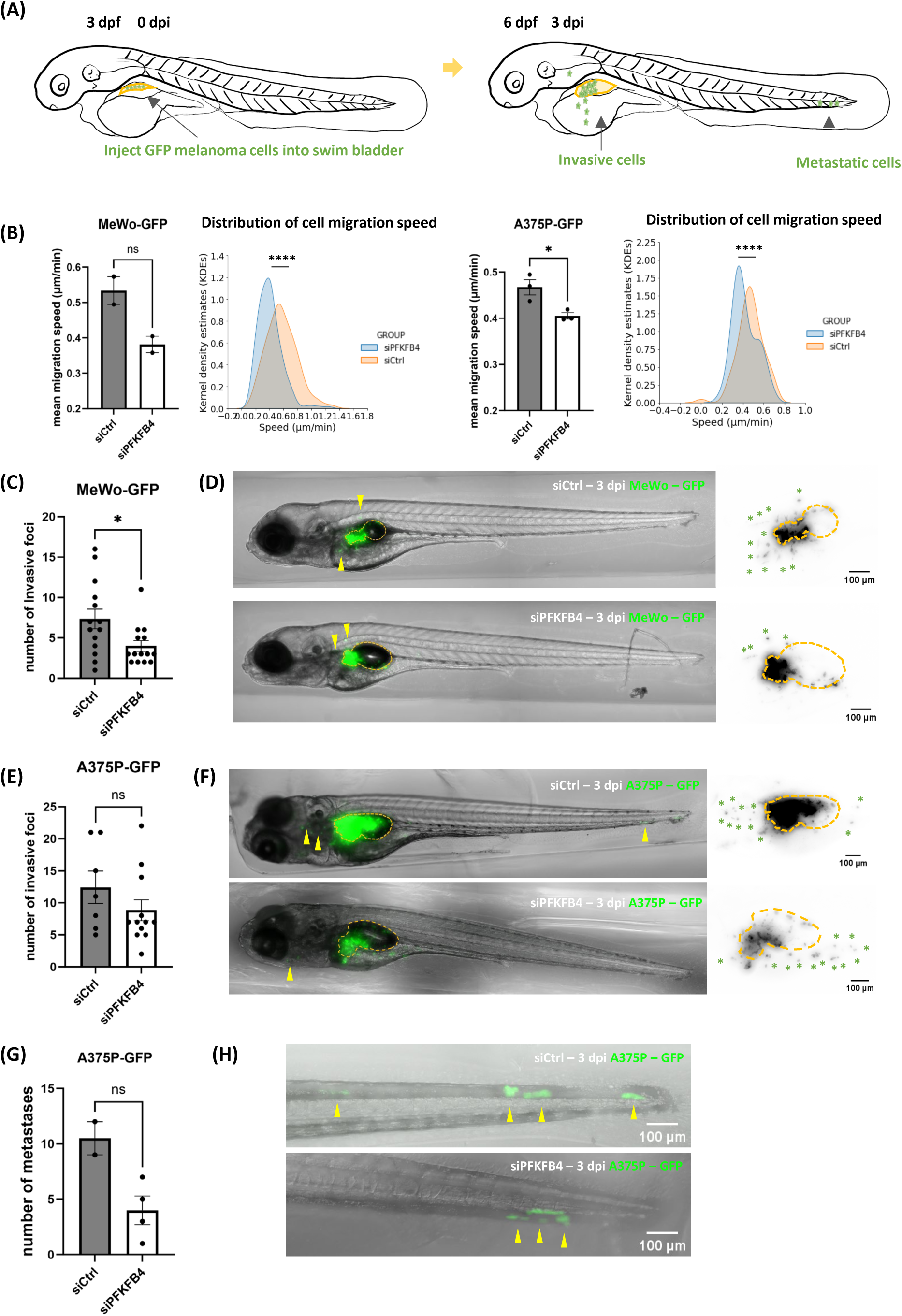
PFKFB4 regulates melanoma cell invasion in the zebrafish xenograft model. **(A)** Schematic representation of the zebrafish xenograft assay. **(B)** Migration capacity *was assessed* in MeWo-GFP or A375P-GFP cells. Starting 48h after siRNA transfection, cell migration was tracked by live microscopy for 16 h. Migration velocity was reduced in both cell lines following PFKFB4 knockdown with siRNA targeting PFKFB4 (siPFKFB4), compared with cells transfected with non-targeting control siRNA (siCtrl). Bar plots represent mean ± SEM of three independent experiments (unpaired *t-*test). Kernel density plots illustrate the distribution of migration speeds in siCtrl versus siPFKFB4 groups. **(C) (E)** Number of invasive MeWo-GFP foci **(C)** or A375P-GFP foci **(E)** per fish at 3 days post injection (dpi) after transfection with siCtrl or siPFKFB4. **(D) (F)** Representative zebrafish larvae at 3 dpi xenografted with MeWo-GFP **(D)** or A375P-GFP **(F)** cells after transfection with siCtrl or siPFKFB4. Asterisks indicate invasive foci detached from the primary tumor. Orange dotted lines mark the swim bladder boundary. **(G)** Number of distant metastatic A375P-GFP foci, defined as cells migrating > 300 μm from the tumor boundary. **(H)** Magnification of the tail region showing A375P-GFP cells metastasizing to caudal hematopoietic tissue. In panels **C**, **E**, and **G**, each dot represents one fish. Statistical significance was determined by unpaired *t-*test. n.s., not significant (P > 0.05); *P < 0.05; dpi, days post injection; dpf, days post fertilization.

Tumor cells that migrate beyond the swim bladder must traverse at least one epithelial layer of the bladder wall, thereby recapitulating key features of cancer invasion. We first evaluated the penetrance of this phenotype. Among larvae injected with MeWo cells, invasive foci were observed in 82% (14/17) of the siCtrl group and 93% (14/15) of the siPFKFB4 group. Similarly, invasive cells were detected in 100% (7/7) of the siCtrl-A375P group and 86% (12/14) of the siPFKFB4-A375P group. Thus, in both models, invasion was highly penetrant, and the penetrance of the phenotype was largely independent of PFKFB4 levels. However, penetrance alone does not capture the extent of the invasive process. We therefore quantified the intensity of invasion by measuring the number of tumor cell foci (clusters of one or a few cells) located outside the swim bladder at 3 dpi. In MeWo-GFP xenografts, invasive foci were observed in tissues surrounding the swim bladder, including liver, pancreas, and yolk sac. The number of invasive foci per larva was significantly reduced following PFKFB4 depletion (n=14) compared to siCtrl group (n=14) (**Figure 2. C, D**). By contrast, in A375P-GFP xenografts, PFKFB4 knockdown did not significantly affect the number of invasive foci (**Figure 2. E, F**).

We next quantified distant metastases by detecting GFP-labelled cells in the larval caudal hematopoietic tissue or head region, located more than 300μm from the swim bladder. In MeWo-GFP xenografts, no metastases were observed regardless of whether the injected cells were transfected with control siRNA or siRNA targeting PFKFB4. By contrast, A375P-GFP cells metastatic dissemination to both sites, with similar frequencies in control cells (29%, n=2/7) and PFKFB4-depleted cells (29%, n=4/14). Among larvae that developed metastases (2/7 siCtrl and 4/14 siPFKFB4), the number of metastatic foci per larvae showed a tendency to decrease after PFKFB4 knockdown, but the difference was not statistically significant (**Figure 2. G, H**). Although the intensity of the metastatic phenotype appeared lower in siPFKFB4 A375P xenografts, the low penetrance of metastasis in this group prevents firm conclusions.

Finally, we reconstructed tumors in 3D from z-stacks and compared primary tumor volumes between control and PFKFB4-depleted conditions. In MeWo-GFP xenografts, no significant difference in tumor volume was observed between control and PFKFB4 knockdown groups (n = 16 and n = 15). In contrast, A375P-GFP tumor volume was significantly reduced following PFKFB4 depletion (n = 7 and n = 8; Figure S2. E, F).

Taken together, these findings highlight PFKFB4 as a context-dependent driver of melanoma growth and invasion *in vivo*, with its impact varying across cell lines.

### PFKFB4 regulates SNAIL2 expression in a melanoma cell line-dependent manner

Previously, we showed that PFKFB4 regulated AKT signaling^37,50,51^, an upstream pathway controlling EMT-TFs such as SNAIL1, SNAIL2 and TWIST^64^. We therefore investigated whether PFKFB4 could influence melanoma cell migration by modulating EMT-TFs expression. Given the cell line-specific effects of PFKFB4 depletion observed *in vivo* in zebrafish, we hypothesized that EMT-TF regulation might also be context-dependent. We found that PFKFB4 and EMT-TFs were expressed at varying levels across melanoma cell lines (**Figure 3. A, Figure S3**). MeWo and SK28 cells strongly expressed SNAIL2, and ZEB2, but showed low SNAIL1 and TWIST expression, and no detectable ZEB1 expression, consistent with a proliferative phenotype. In contrast, A375M cells expressed high levels of SNAIL1, TWIST and ZEB1, but low SNAIL2 and ZEB2, consistent with an invasive phenotype. A375P cells displayed a similar profile to A375M, with strong ZEB1 expression, no detectable ZEB2, and lower levels of other EMT-TFs. These heterogenous EMT-TF expression patterns and their association with melanoma phenotype switching were consistent with previous reports^18^. Importantly, although all tested cell lines exhibited reduced motility upon PFKFB4 depletion *in vitro*^37^, their diverse EMT-TF expression profiles suggests that PFKFB4 may influence cell motility either independently of EMT-TFs, via distinct EMT-TFs in different cell lines, or through a combination of both mechanisms.

**Figure 3.**
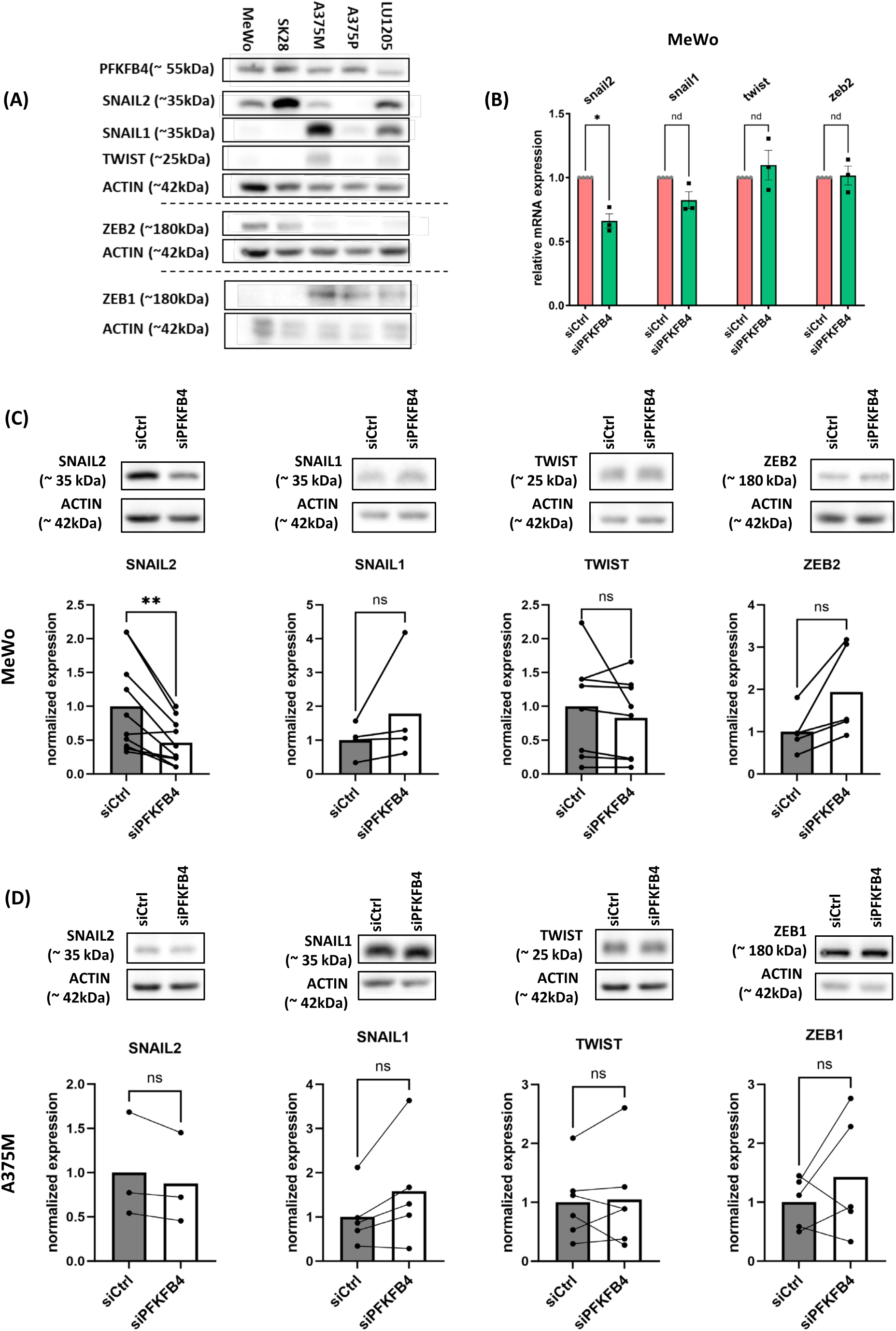
PFKFB4 regulates SNAIL2 expression in a melanoma cell line-dependent manner. **(A)** Protein expression levels of the EMT-TFs SNAIL2, SNAIL1, TWIST, ZEB2 and ZEB1 across different melanoma cell lines. **(B)** mRNA expression of EMT-TFs in MeWo cells assessed by qPCR 48h after transfection with siCtrl or siPFKFB4. Statistical analysis was performed using an unpaired t-test. **(C, D)** Normalized protein expression of EMT-TFs in MeWo **(C)** or A375M **(D)** cells, 48h after transfection with siCtrl or siPFKFB4. Each dot represents one biological replicate with paired values connected by lines. Statistical analysis was performed using a paired *t-*test. n.s, not significant (P > 0.05); *P < 0.05; **P < 0.01.

We next examined EMT-TF expression following PFKFB4 depletion. In MeWo cells, both SNAIL2 mRNA and protein levels were reduced by about half after PFKFB4 knockdown (detected by by qPCR, n=3; and Western blot, n=10) (**Figure 3. B, C, Figure S4. A**), whereas other EMT-TFs remained unchanged. By contrast, no alteration in EMT-TF expression was observed in A375M and A375P cell lines depleted for PFKFB4 (**Figure 3. D, Figure S4. A, B**). These findings indicated that EMT-TFs regulation by PFKFB4 was cell line-dependent and largely restricted to SNAIL2. Since PFKFB4 depletion reduced motility in MeWo and A375 cell lines but downregulated SNAIL2 only in MeWo, this suggested that PFKFB4 might regulate motility via SNAIL2 in MeWo, whereas in A375 cell lines its effect on motility was more likely independent of EMT-TFs. This observation prompted us to investigate whether a PFKFB4-SNAIL2 axis contributed to MeWo cell motility.

### PFKFB4 regulation of MeWo cell migration is independent of SNAIL2

To determine whether the reduction of SNAIL2 contributed to the decreased MeWo cell migration observed after PFKFB4 depletion, we first depleted SNAIL2 expression by siRNA transfection. Cell velocity *in vitro* was significantly reduced compared with cells transfected with a control siRNA, confirming that SNAIL2 was important for MeWo melanoma cell migration (**Figure 4. A**). We then performed rescue experiments to test a potential epistatic relationship between PFKFB4 and SNAIL2. Co-transfection of human PFKFB4 siRNA with a plasmid encoding the *Xenopus laevis Snail2* gene (*x-Snail2*, 89% homologous to human *SNAIL2*) did not restore migration in PFKFB4-depleted cells (**Figure 4. B, C**). By contrast, expression of Xenopus laevis PFKFB4 (95% homologous to the human PFKFB4 protein and insensitive to the siRNA) successfully rescued the migration defect but did not reverse the decrease in SNAIL2 levels (**Figure 4. D, E, F**). Together these results indicate that PFKFB4 promotes MeWo cell migration through mechanisms that are uncoupled from its regulation of SNAIL2 expression.

**Figure 4.**
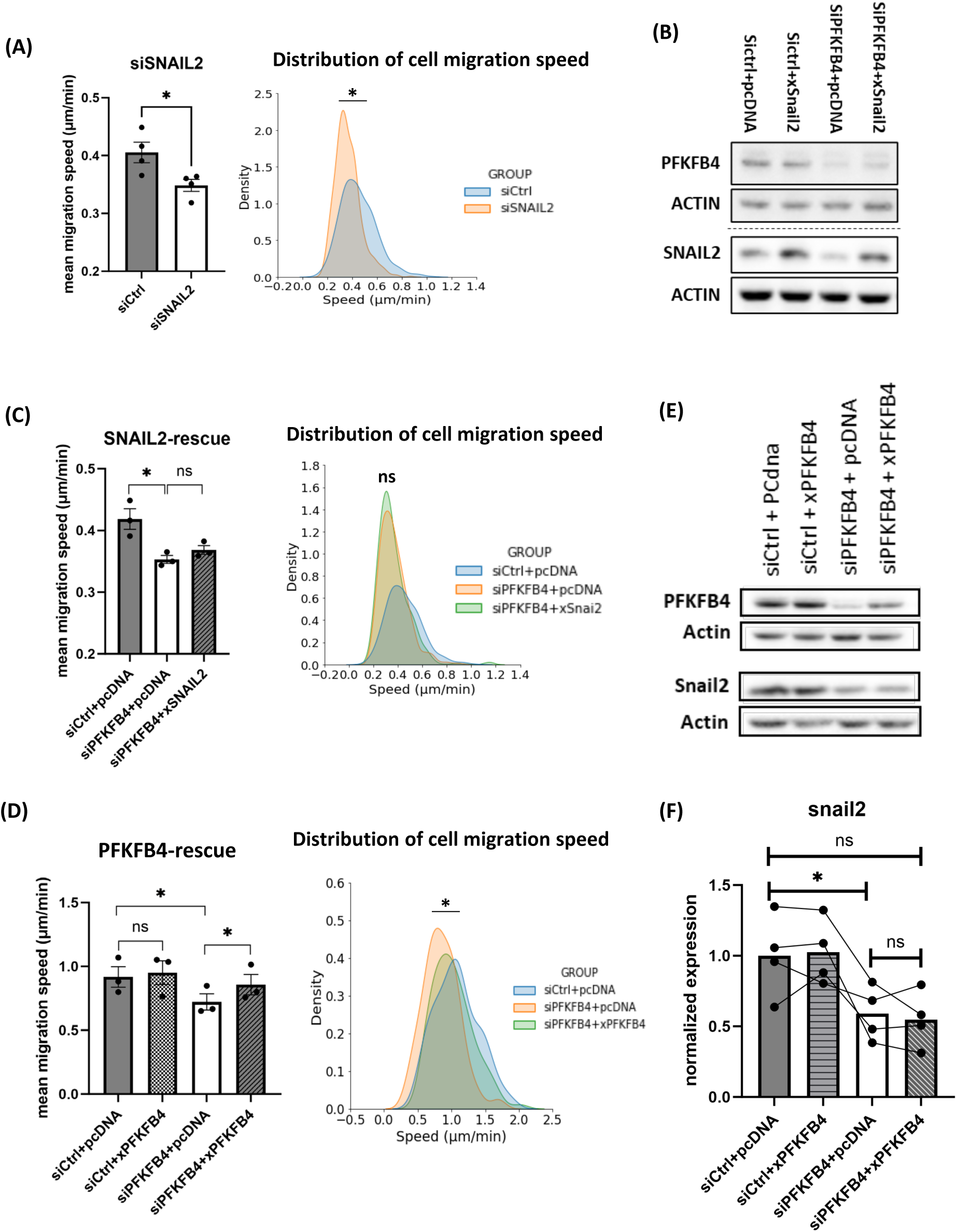
PFKFB4 regulates MeWo cell migration independently of SNAIL2. **(A)** Mean migration velocity of MeWo cells 48 h after transfection with siCtrl or siRNA targeting SNAIL2 (siSNAIL2). Kernel density plots illustrate the distribution of migration speeds in control versus SNAIL2 knockdown groups. **(B,C)** Expression of PFKFB4 and SNAIL2 proteins **(B)** and mean migration velocity **(C)** of MeWo cells transfected with siCtrl or siRNA targeting PFKFB4 (siPFKFB4), combined with empty pcDNA vector or a plasmid expressing *Xenopus laevis* SNAIL2 (xSNAIL2). Kernel density plots show the distribution of migration speeds across groups. xSNAIL2 failed to rescue the migration defect induced by PFKFB4 knockdown. **(D)** Mean migration velocity of MeWo cells after siCtrl or siPFKFB4 transfection, combined with empty pcDNA vector or a plasmid expressing Xenopus laevis PFKFB4 (xPFKFB4). Kernel density plots show the distribution of migration speeds. xPFKFB4 was able to rescue the migration defect caused by PFKFB4 knockdown. **(E, F)** Protein expression levels of PFKFB4 and SNAIL2 **(E)** and quantification from four biological replicates **(F)** in MeWo cells transfected with siCtrl or siPFKFB4, combined with empty pcDNA or xPFKFB4 plasmid. xPFKFB4 failed to rescue the decrease in SNAIL2 protein levels caused by PFKFB4 knockdown. In panels **A**, **C**, and **D**, bar plots represent mean ±SEM and were analyzed using an unpaired *t-*test. **(F)** Each dot represents one biological replicate, with paired values connected by lines; significance was assessed using a paired *t-*test. n.s., not significant (P > 0.05); *P < 0.05.

## DISCUSSION

Previously, we showed that PFKFB4 positively regulated cell migration *in vitro* in various melanoma cell lines, including MeWo and A375M^37^. Moreover, high PFKFB4 expression is associated with poor patient survival, suggesting that PFKFB4 may contribute to melanoma progression (Figure 1). Here, we used a larval zebrafish xenograft model, injecting human melanoma cell lines into the swim bladder, to evaluate the effects of PFKFB4 on tumor growth, invasion, and metastasis *in vivo* (Figure 2), and to probe whether EMT-TFs such as SNAIL2 could participate in these processes *in vitro* (Figures 3, 4).

Both MeWo and A375P cell lines are derived from human metastatic cutaneous melanoma. In nude mice, MeWo cells are highly invasive and metastatic^65^, while A375P is a poorly metastatic subclone of the A375 lineage^62,63^. In the zebrafish xenograft model, however, both cell lines displayed invasive behavior, but only A375P cells generated distant metastases, highlighting a model-dependent variability in phenotype.

In MeWo xenografts, PFKFB4 depletion had no effect on primary tumor growth or on the penetrance of the invasive phenotype, which remained high in both siCtrl and siPFKFB4 groups. However, the intensity of invasion was significantly reduced, as fewer invasive foci were detected in surrounding tissues. No distant metastases were observed in either siCtrl or siPFKFB4 MeWo xenografts, precluding evaluation of a role for PFKFB4 in metastatic dissemination in this line.

In contrast, in A375P xenografts, PFKFB4 depletion significantly reduced primary tumor volume but had no significant effect on either the penetrance or intensity of invasion, both of which remained high. Similarly, PFKFB4 knockdown did not significantly alter the penetrance or intensity of the metastatic phenotype, which was intrinsically low in this line. Although reduced tumor size could theoretically limit the pool of disseminating cells, this explanation seems unlikely since invasive foci were not decreased despite smaller tumors. Taken together, these results indicate that PFKFB4 regulates MeWo cell invasion *in vivo* but has no significant effect on invasion or metastasis in A375P cells, while impacting tumor growth only in the latter. Thus, PFKFB4 influences melanoma progression in a strongly cell line–dependent manner.

To interpret this variability, we considered both the genotype and cellular characteristics of each melanoma cell line. In addition to their BRAF or NRAS mutational status, melanoma cell lines can be broadly categorized into “proliferative” (e.g. MeWo) or “invasive” (e.g. A375) states based on their EMT-TF expression profiles and specific functional signatures^21,55^. The BRAF^V600E^ mutation activates MAPK/ERK signaling and has been associated with phenotype switching from a proliferative (SNAIL2^high^, ZEB2^high^, ZEB1^low^, TWIST1^low^) to an invasive (SNAIL2^low^, ZEB2^low^, ZEB1^high^, TWIST1^high^) state^21^. EMT-TF expression patterns in MeWo (BRAF-WT, SNAIL2^high/^ZEB2^high^) and A375 (BRAF^V600E^, ZEB1^high^/TWIST1^high^) were broadly consistent with this framework, although the relationship between BRAF genotype and EMT status was not strictly linear (Figure S3). Our xenograft data also provide partial support for this classification: MeWo tumors, although smaller on average, displayed greater growth (higher D3/D0 volume ratio; Figure S2. E), whereas A375P tumors, produced a higher number of invasive foci (Figure 2. C & E). These results may reflect, at least in part, the proliferative status of MeWo cells and the invasive status of A375P cells. Importantly, PFKFB4 depletion differentially impacted these behaviors, reducing invasion in MeWo cells but primarily affecting primary tumor growth in A375P cells.

EMT-TFs are regulated by the AKT-GSK3 signaling axis^52–54^ which itself has been shown to depend on PFKFB4^37,50,51^. Using MeWo and A375M cell lines, we found that PFKFB4 did not affect the expression of SNAIL1, ZEB1, ZEB2 or TWIST. Instead, PFKFB4 specifically regulated SNAIL2 expression in MeWo cells, but not in A375 cells (which are SNAIL2^low^). Although, SNAIL2 downregulation impaired MeWo cell migration *in vitro*, our rescue experiments suggested that this effect was not epistatically linked to PFKFB4, suggesting that PFKFB4 and SNAIL2 contribute to MeWo cell migration through partially independent pathways. Whether the reduction of SNAIL2 expression upon PFKFB4 loss is mediated by AKT-dependent mechanisms remains to be determined.

In summary, our study demonstrates that PFKFB4 regulates melanoma progression in a cell line–dependent manner. In zebrafish xenografts, PFKFB4 depletion reduced invasion in MeWo cells without affecting tumor growth, whereas in A375P cells it decreased primary tumor growth but had no significant effect on invasion or metastasis. These results suggest that PFKFB4 may preferentially contribute to the limiting trait of a given cell line — invasion in proliferative MeWo cells and proliferation in invasive A375P cells. Although speculative, this possibility warrants further testing across additional melanoma lines to determine whether PFKFB4 consistently uncouples proliferation and invasion from the classical phenotype classification. Mechanistically, PFKFB4 selectively regulated SNAIL2 expression in MeWo cells, but this effect was not epistatically linked to its role in migration, suggesting that alternative or parallel pathways are involved. Given the association between high PFKFB4 expression and poor patient prognosis, further studies are warranted to dissect its mechanisms of action in melanoma dissemination and to evaluate its potential as a therapeutic target.

## ACKNOWLEDGEMENTS

We thank Drs L. Larue, N. Manel, G. Seano and C. Janke for kind gift of cells and plasmids. We are grateful to Drs J.-F. Riou, E. Theveneau and S. Druillennec for insightful scientific discussions and to L. Besse and M.-N. Soler for their help on live-imaging (CurieCoreTech Multimodal Imaging Center (UAR2016 / US43)). This work was funded by grants from Agence Nationale pour la Recherche (ANR-21-CE13-0028), Institut Universitaire de France and Fondation pour la Recherche Medicale (FDT202404018665) to AHMB; and by European Union Horizon 2020 Marie Skłodowska-Curie grant No 860635, NEUcrest ITN to AHMB and KK.

## SUPPLEMENTARY FIGURES

**Figure S1. The expression of PFKFB4 in human melanoma patients**

Box plots showing PFKFB4 mRNA expression levels in normal skin, nevi, melanoma in situ, primary melanoma, and metastatic melanoma. Data were extracted from the GSE7553 (**A**), GSE3189 (**B**), and GSE8401 (**C**) datasets. Statistical significance was assessed using one-way ANOVA.

**Figure S2. Tumor growth in the zebrafish xenograft model**

**(A)** Morphology of MeWo-GFP cells compared with parental MeWo cells. **(B)** PFKFB4 expression in MeWo-GFP cells and A375P-GFP cells 48 h after transfection with non-targeting control siRNA (siCtrl) or PFKFB4-specific siRNA (siPFKFB4). **(C)** The penetrance of invasive or metastatic phenotypes in the zebrafish xenograft model. **(D)** Representative zebrafish larvae at the day of injection (D0). **(E-F)** Tumor growth ratios at three days post injection (dpi) relative to D0 in zebrafish xenografted with MeWo-GFP **(E)** or A375P-GFP **(F)** cells. Each dot represents one fish. Statistical significance was assessed using an unpaired t-test; n.s. not significant (P > 0.05); **P < 0.01.

**Figure S3. mRNA expression of PFKFB4 and EMT-TFs in human melanoma cell lines.**

Heatmaps show the mRNA expression of EMT-TFs, PFKFB4, CDH1, and CDH2 across multiple human melanoma cell lines. Data were extracted from Rambow, F. et al., 2015^55^.

**Figure S4. PFKFB4 does not affect EMT-TFs expression in A375P cells.**

**(B)** PFKFB4 expression in MeWo cells or A375M cells 48 h after transfection with non-targeting control siRNA (siCtrl) or PFKFB4-specific siRNA (siPFKFB4). **(B)** Protein levels of SNAIL1, TWIST, and ZEB1 were not significantly altered after PFKFB4 knockdown in A375P-GFP cells.

## Notes

### Competing Interest Statement

The authors have declared no competing interest.

