## Supplementary figures for "Zebrafish Xenografts Reveal a Context-dependent Role of PFKFB4 in Melanoma Cell"

**Figure S1. The expression of PFKFB4 in human melanoma patients**

Box plots showing PFKFB4 mRNA expression levels in normal skin, nevi, melanoma in situ, primary melanoma, and metastatic melanoma. Data were extracted from the GSE7553 (**A**), GSE3189 (**B**), and GSE8401 (**C**) datasets. Statistical significance was assessed using one-way ANOVA.

**Figure S2. Tumor growth in the zebrafish xenograft model**

**(A)** Morphology of MeWo-GFP cells compared with parental MeWo cells. **(B)** PFKFB4 expression in MeWo-GFP cells and A375P-GFP cells 48 h after transfection with non-targeting control siRNA (siCtrl) or PFKFB4-specific siRNA (siPFKFB4). **(C)** The penetrance of invasive or metastatic phenotypes in the zebrafish xenograft model. **(D)** Representative zebrafish larvae at the day of injection (D0). **(E-F)** Tumor growth ratios at three days post injection (dpi) relative to D0 in zebrafish xenografted with MeWo-GFP **(E)** or A375P-GFP **(F)** cells. Each dot represents one fish. Statistical significance was assessed using an unpaired t-test; n.s. not significant ($P > 0.05$); $^{**}P < 0.01$.

**Figure S3. mRNA expression of PFKFB4 and EMT-TFs in human melanoma cell lines.**

Heatmaps show the mRNA expression of EMT-TFs, PFKFB4, CDH1, and CDH2 across multiple human melanoma cell lines. Data were extracted from Rambow, F. et al., 2015[55].

**Figure S4. PFKFB4 does not affect EMT-TFs expression in A375P cells.**

**(A)** PFKFB4 expression in MeWo cells or A375M cells 48 h after transfection with non-targeting control siRNA (siCtrl) or PFKFB4-specific siRNA (siPFKFB4). **(B)** Protein levels of SNAIL1, TWIST, and ZEB1 were not significantly altered after PFKFB4 knockdown in A375P-GFP cells.

**Zhou et al.,**
**Figure S1.**

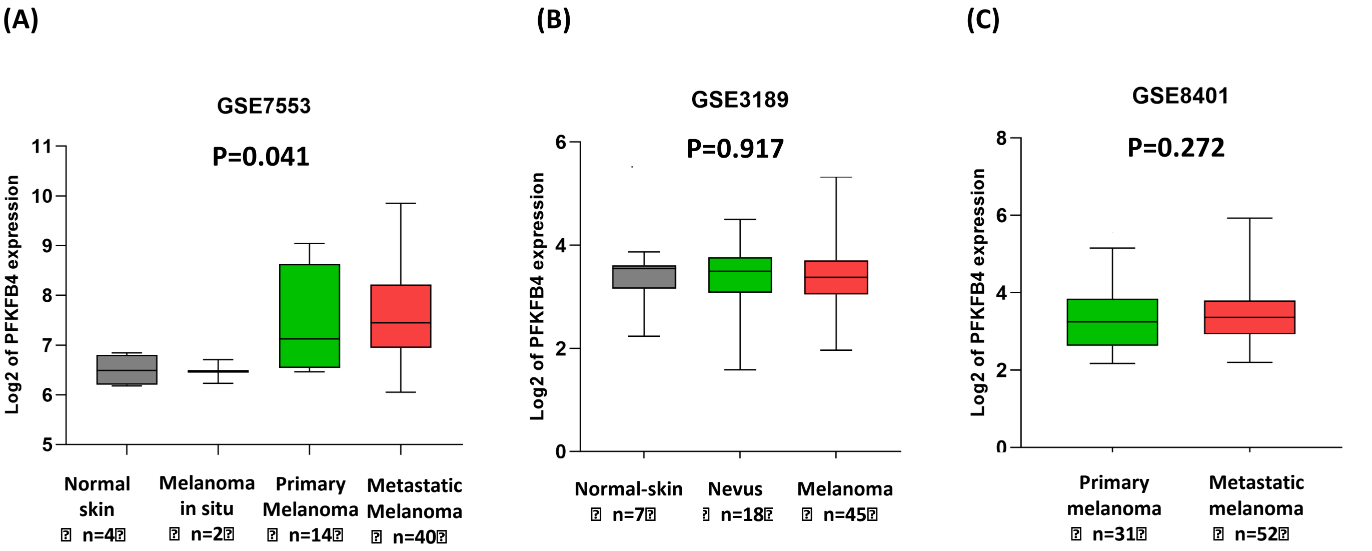

**Zhou et al.,**
**Figure S2.**

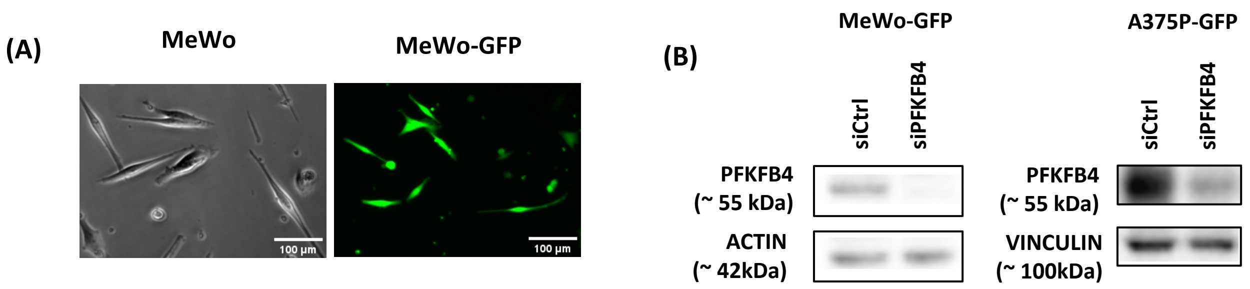


(C)

| The penetrance of phenotypes | | | | |
|---|---|---|---|---|
| | Mewo | | A375P | |
| | siCtrl | siPFKFB4 | siCtrl | siPFKFB4 |
| # larva included | 17 | 15 | 7 | 14 |
| # larva with invasive phenotype | 14 | 14 | 7 | 12 |
| # larva with metastasis phenotype | 0 | 0 | 2 | 4 |

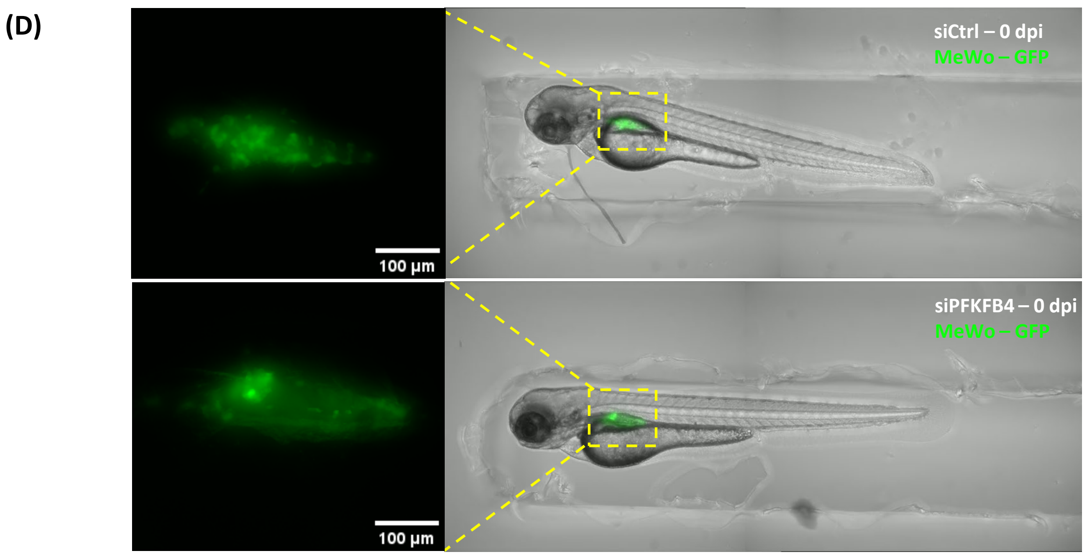


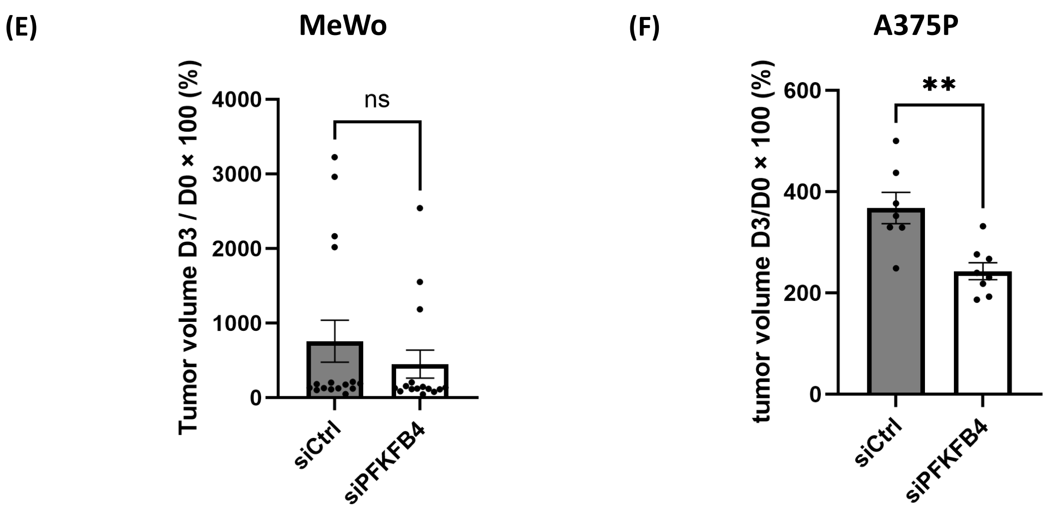

**Zhou et al.,**
**Figure S3.**

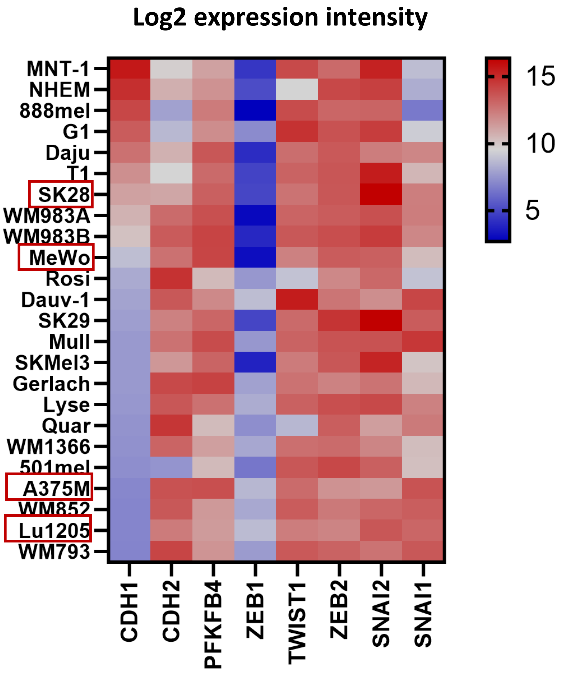

**Zhou et al.,**
**Figure S4.**

(A)

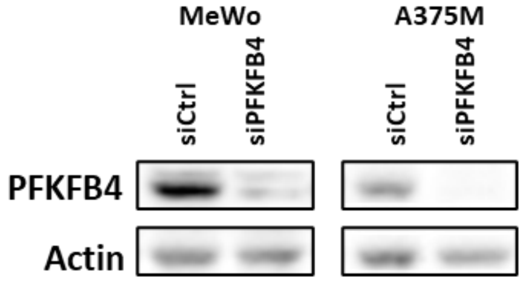


(B)

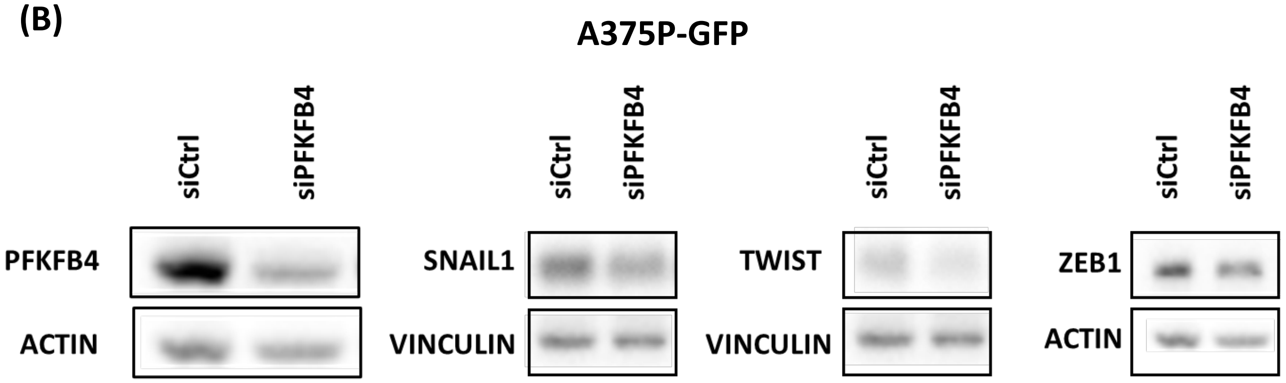
